# Strategies to improve photosynthetic nitrogen-use efficiency with no yield penalty: lessons from late-sown winter wheat

**DOI:** 10.1101/379552

**Authors:** Lijun Yin, Haicheng Xu, Shuxin Dong, Jinpeng Chu, Xinglong Dai, Mingrong He

## Abstract

**Highlight:** Optimal *N* allocation at several integration levels accounts for improved canopy *PNUE* while maintaining high grain yield in winter wheat

**Abstract:** Improving canopy photosynthetic nitrogen-use efficiency (*PNUE*) may maintain or even increase yield with reduced *N* input. In this study, later-sown winter wheat was studied to reveal the mechanism underlying improved canopy *PNUE* while maintaining high yield. *N* allocation at several levels was optimised in late-sown wheat plants. *N* content per plant increased. Increased *N* was allocated to the flag leaf and second leaf, and to ribulose-1, 5-bisphosphate carboxylase/oxygenase (*Rubisco*) in upper leaves. Constant or reduced *N* was allocated to leaf 3, leaf 4, and *Rubisco* in lower leaves. The specific green leaf area nitrogen (*SLN*) of upper leaves increased, while that of lower leaves remained unchanged or decreased. *N* allocation to the cell wall decreased in all leaves. As a result, the maximum carboxylation rate of upper leaves increased, and that of lower leaves remained constant or decreased. CO_2_ diffusion capacity was enhanced in all leaves. Outperformance by light-saturated net photosynthetic rate (*P_max_*) over *SLN* led to improved *PNUE* in upper leaves. Enhanced *Pmax* coupled with unchanged or decreased *SLN* resulted in improved *PNUE* in lower leaves. High yield was maintained because enhanced photosynthetic capacity at the leaf and whole plant levels compensated for reduced canopy leaf area.

## Introduction

Wheat (*Triticum aestivum* L.) provides 20% of the calories and protein consumed by humans (Reynolds *et al.*, 2012). An increase in crop yield by 70% is needed if we are to meet the projected demand for food by 2050 (Tilman *et al.*, 2011; Ray *et al.*, 2013). The amount of nitrogen (*N*) applied will increase with the growing demand for food production in the future (Li *et al.*, 2017). Increased economic costs and environmental concerns have heightened the desire to reduce crop *N* input while maintaining or even increasing grain yield (Cassman *et al.*, 2003; Davidson *et al.*, 2015; Zhang *et al.*, 2015). Therefore, improving N-use efficiency (*NUE*) has become a top priority for crop improvement. *NUE* is defined as grain yield per unit of *N* available (from soil and/or fertiliser) and can be further divided into N-uptake efficiency and N-utilisation efficiency (*UTE*) (Moll *et al.*, 1982). *UTE*, defined as grain yield per unit of *N* taken up, is an important parameter for determining the efficiency with which crop plants utilise *N* to achieve growth and grain yield (Foulkes *et al.*, 2009).

At the end of the 1970s, the concept of plant *N* productivity, defined as the increase in plant dry matter per unit time and per unit *N* content, was introduced to interpret the dependency of plant growth on internal *N* (Ingestad *et al.*, 1979). Following Lambers *et al.* (1990) and Garnier *et al.* (1995), plant *N* productivity was expressed as the product of *N* allocation to leaves within the plant and photosynthetic *N* use efficiency (*PNUE).* The latter was defined as the ratio between photosynthetic rate and *N* concentration in leaves. As most of the grain dry matter at maturity in wheat is contributed by photosynthates produced by leaves during the post-anthesis stage (Roberto *et al.*, 2010; Carmo-Silva *et al.*, 2017), *UTE* at the whole-plant level is dependent on *N* allocation to leaves and the *PNUE* of leaves during the post-anthesis stage.

Plants change *N* allocation to maximise their carbon assimilation at several integration levels. First, they allocate a given amount of *N* over a small or a large plant population through trade-offs between plant density per unit of land and *N* content in individual plants. Second, they change the fraction of *N* invested in leaves, stems and roots. Third, they modulate leaf area per unit *N* invested in leaves by altering their anatomy. Fourth, they change the relative investment of *N* among photosynthetic components. Small changes in *N* allocation can greatly affect the light-saturated photosynthetic rate (*P_max_*) and *PNUE*, and therefore plant performance (Feng *et al.*, 2009).

Strategies to improve *PNUE* have been proposed for many species (Poorter *et al.*, 1998; Davey *et al.*, 1999; Pang *et al.*, 2014; Rotundo *et al.*, 2016). Most studies have proposed that the potential benefit of increased photosynthetic capacity for *PNUE* can be realised only when it is not associated with increases in leaf mass per area (*LMA*, g m^−2^) or specific leaf *N* content (N content per unit leaf area, *SLN*), as an increase in *LMA* positively affects *SLN* and therefore reduces *PNUE* (Field and Mooney, 1986; Hirose *et al.*, 1994; Hikosaka *et al.*, 1995; Boote *et al.*, 2003; Anand *et al.*, 2007). However, a strategy that results in lower *SLN* may limit crop yield under some conditions, particularly those designed to produce high yields. Actually, previous studies have suggested that *N* remobilisation from vegetative tissues may be essential as a mobilisable *N* reservoir to sustain grain yield in cereal crops (Horton, 2000; Barbottin *et al.*, 2005). The amount of *N* accumulated at anthesis largely determines the amount of *N* remobilised during grain filling (Martre *et al.*, 2003; Pask *et al.*, 2012). Taken together, these findings suggest that alternative approaches need to be explored to improve *PNUE* while maintaining high or even increasing grain yield.

Interspecific or intraspecific variations in *PNUE* have been explained by differences in the fraction of light absorbed by the leaf, CO_2_ partial pressure at the intercellular space or at carboxylation sites within chloroplasts, *N* allocation to photosynthetic versus non-photosynthetic functions, *N* partitioning between light harvesting complexes, electron transport and CO_2_ fixation, activation state or specific activity of ribulose-1, 5-bisphosphate carboxylase/oxygenase (*Rubisco*), respiration in the light, and *SLN* (Field and Mooney, 1986; Evans *et al.*, 1989; Quick *et al.*, 1991; Lambers *et al.*, 1992; Pons *et al.*, 1994; Zhu *et al.*, 2007).

Global warming over past decades has provided an additional growing period prior to wintering that has encouraged farmers to delay the winter wheat sowing date (Xiao *et al.*, 2013, 2015). Previous studies have indirectly suggested that delayed sowing of winter wheat may have advantages in crop productivity as a function of plant *N* use (Widdowson *et al.*, 1987; Ehdaie *et al.*, 2001; Weiss *et al.*, 2003; Sun *et al.*, 2007; Jalota *et al.*, 2013; Ding *et al.*, 2016; Rasmussen *et al.*, 2016). Our recent study suggested that delayed sowing improves *UTE* while maintaining a high yield by increasing spike grain weight with fewer spikes per unit area (Yin *et al.*, 2018), suggesting concurrent improvement in *PNUE* and grain productivity at the whole-plant level. The following questions have arisen from these results: (i) What causes late-sown wheat plants to have a higher *PNUE?* (□) How is coupling between improvement in *PNUE* at the whole-plant level and high grain yield at the canopy level achieved? To answer these questions, photosynthetic traits in plants, such as leaf gas exchange, chlorophyll fluorescence, *Rubisco* catalytic properties, and CO_2_ diffusion capacity, were investigated along with *N* allocation at the canopy, whole-plant, leaf, and cellular levels. Our main goal was to test the hypothesis that optimal *N* allocation at several integration levels improves *PNUE* and grain productivity at the whole-plant level, which in turn results in improved canopy *PNUE* while maintaining high yield.

## Materials and methods

### Plant material and growing conditions

Tainong 18, a widely planted winter wheat cultivar, was grown in the field at the experimental station of Shandong Agricultural University, Taian, Shandong, China during the 2015–2016 and 2016–2017 growing seasons. The preceding crop was summer maize. The soil was sandy loam with a pH of 8.0. The contents of organic matter (Walkley and Black method), total *N* (semi-micro Kjeldahl method), available phosphorus (*P*; Olsen method), and available potassium (*K*; Dirks–Sheffer method) in the 0–20-cm soil layer were 12.0, 1.0, 25.1, and 47.0 mg kg^−1^ during 2015–2016 and 12.1, 1.0, 25.3, and 47.1 mg kg^−1^ during 2016–2017, respectively. Rainfall levels during the growing seasons of 2015–2016 and 2016–2017 were 144.9 and 168.3 mm, respectively.

Seeds were sown at a density of 405 plants m^−2^, the optimal planting density of Tainong 18 for higher yield and *NUE* (Dai *et al.*, 2013), in 2015 and 2016 on 8 October (normal sowing) and 22 October (late sowing) using a 12-row planter with 0.25-m row spacing. The cumulative temperature values (sum of daily average air temperature) prior to wintering of the normal and late-sown treatments were 679.4 and 444.5°C d during the 2015–2016 growing season and 682.4 and 449.5°C d during the 2016–2017 growing season, respectively. The plots were arranged in a completely random design with three replicates. The size of each subplot was 20.0 × 3.0 m. Basal fertilisation of each subplot included *N* as urea, *P* as calcium superphosphate, and *K* as potassium chloride at rates of 120 kg ha^−1^ *N*, 80 kg ha^−1^ P2O5, and 120 kg ha^−1^ K2O, respectively. An additional 120 kg ha^−1^ *N* as urea was applied at the beginning of the jointing stage. Irrigation was carried out before wintering, at jointing, and at anthesis, with approximately 60 mm each time. Pests and diseases were controlled chemically. No significant incidences of pests, diseases, or weeds occurred in any of the subplots.

### Crop measurement Biomass and nitrogen content of individual plant and leaves

Plants on 0.2 m^2^ were taken as samples and counted in each subplot at 7-day intervals from anthesis to maturity. All individual plants were divided into flag leaf, second leaf, leaf 3, leaf 4, and the remaining parts. The planar green area of each leaf was measured (in cm^−2^) using a green area meter (Li-Cor 3100, Li-Cor, Inc., Lincoln, NE, USA). Biomass was measured after oven drying to constant mass at 75°C. The samples were ground, and *N* mass per unit dry mass was determined using an elemental analyser (Rapid *N* Exceed, Elementar, Langenselbold, Germany). The *LMA* (g m^−2^ green leaf area) and *SLN* (g *N* m^−2^ green leaf area) values were calculated.

### Grain yield, yield components, plant *N* productivity, and UTE

Plants were harvested from a 2.0-m × 6-row (1.5 m) quadrat in each subplot as described by Dai *et al.* (2013). The grain was air-dried, weighed, and adjusted to standard 12% moisture content (88% dry matter, kg ha^−1^). This was considered grain dry matter yield.

Plant *N* productivity was defined as the increase in plant dry matter per unit time and per unit *N* content. *UTE* was defined as grain yield per unit of *N* taken up (Moll *et al.*, 1982).

### Biomass and nitrogen content of the cell wall

Biomass and nitrogen content of the cell wall were measured according to the procedures described by Lamport (1965) and Onoda *et al.* (2004). Approximately 10 mg of freeze-dried leaves was extracted in 1.5 mL of buffer (50 mm tricine, pH 8.1) containing 1% PVP40 (average molecular weight 40,000, product no. 1407; Sigma Chemical Co., St Louis, MO, USA). The sample was vortexed and centrifuged at 12,000 *g* for 5 min (AG 5424; Eppendorf, Hamburg, Germany), and the supernatant was carefully removed. The pellet was resuspended in buffer without PVP containing 1% sodium dodecyl sulphate (SDS), incubated at 90°C for 5 min and centrifuged at 12,000 *g* for 5 min. This was repeated, and then two washes with 0.2 m KOH, two washes with deionised water, and then two washes with ethanol were carried out. The tube containing the pellet was oven-dried at 80°C. The remaining dry mass of the pellet was assumed to represent the leaf cell wall biomass, and *N* content was determined on 2–5 mg of material using the elemental analyser.

### Biomass and Rubisco nitrogen content

The *Rubisco* content of each layer leaf at anthesis was determined according to Makino *et al.* (1985, 1986). Briefly, leaves were sampled and immersed in liquid *N* and then stored at −70°C. A 0.5-g aliquot of leaves was ground in a buffer solution containing 50 mM Tris-HCl (pH 8.0), 5 mM β-mercaptoethanol, and glycerol 12.5% (v/v), and the extracts were centrifuged for 15 min at 1,500 *g* at 2°C. The supernatant was mixed with dissolving solution containing 2% (w/v) SDS, 4% (v/v) β-mercaptoethanol, and 10% (v/v) glycerol, and the mixture was boiled in water for 5 min for the protein electrophoresis assay. An electrophoretic buffer system was used with sodium dodecyl sulphate–polyacrylamide gel electrophoresis in a discontinuous buffer system with a 12.5% (w/v) separating gel and a 4% (w/v) concentrated gel. The gels were washed with deionised water several times, dyed in 0.25% Coomassie Blue staining solution for 12 h, and decolourised until the background was colourless. Large subunits and relevant small subunits were transferred to a 10-ml cuvette with 2 ml of formamide and washed in a 50°C water bath at room temperature for 8 h. The wash solution was measured at 595 nm using background glue as the blank and bovine serum albumin as the standard protein. Because the amount of *N* per unit *Rubisco* is 16% (Field and Mooney, 1986), *Rubisco N* content per unit leaf area was calculated as *Rubisco* content multiplied by 16%.

### Relative amount of mRNA

Total *RNA* was extracted from frozen leaf discs using Trizol according to the manufacturer’s specifications. *RNA* yield was determined using a NanoDrop 2000 spectrophotometer (Thermo Scientific, Waltham, MA, USA), and integrity was evaluated by agarose gel electrophoresis and ethidium bromide staining.

A two-step reaction process of reverse transcription and polymerase chain reaction (PCR) was used for quantification. Each reverse transcription reaction had two steps. The first step was 0.5 μg *RNA*, 2 μl of 4 × g *DNA* wiper Mix and 8 μl of nuclease-free H_2_O. Reactions were performed in a GeneAmp^®^ PCR System 9700 (Applied Biosystems, Foster City, CA, USA) for 2 min at 42°C. The second step was to add 2 μl of 5× HiScript II Q reverse transcription SuperMix IIa. Reactions were performed in a GeneAmp^®^ PCR System 9700 for 10 min at 25°C, 30 min at 50°C, and 5 min at 85°C. The 10 μl of reverse transcription reaction mix was diluted ×10 in nuclease-free water and held at −20°C. Real-time PCR was performed using the LightCycler^®^ 480 II Real-time PCR instrument (Roche, Basel Switzerland) with 10 μl of PCR reaction mixture that included 1 μl of cDNA, 5 μl of 2× QuantiFast^®^ SYBR^®^ Green PCR Master Mix (Qiagen, Hilden, Germany), 0.2 μl of forward primer, 0.2 μl of reverse primer and 3.6 μl of nuclease-free water. Reactions were incubated in a 384-well optical plate (Roche) at 95°C for 5 min, followed by 40 cycles of 95°C for 10 s and 60°C for 30 s. Each sample was run in triplicate for the analysis. At the end of the PCR cycle, a melting curve analysis was performed to validate specific generation of the expected PCR product. The primer sequences were designed in the laboratory and synthesised by Generay Biotech (Generay, PRC) based on the *mRNA* sequences obtained from the NCBI database as follows: AGTAGCTGCCGAATCTTCT.

The expression levels of *mRNAs* were normalised to GAPDH and were calculated using the 2^−ΔΔCt^ method (Livak and Schmittgen, 2001).

### Activated and inactivated Rubisco content

Initial and total *Rubisco* activities were determined according to a procedure described by Keys and Parry (1990). Initial activity was determined by adding 25 μl of supernatant to 475 ml of a CO_2_-free assay buffer containing 100 mM bicine, pH 8.2, and 20 mM MgCl_2_, to which NaH^14^CO_3_ (7.4 kBq μmol^−1^) and *RuBP* had been added to concentrations of 10 and 0.4 mM, respectively, immediately prior to adding the extract. Total activity was determined by incubating 20 ml of extract for 3 min in 980 ml of the same assay buffer without *RuBP*, allowing for carbamylation of all available active sites. The assay was started by adding 0.4 mM *RuBP* as indicated above. The *Rubisco* activation state was determined from the ratio of initial to total activity. The inactive *Rubisco* content was the difference between the total amount of *Rubisco* and active *Rubisco* content.

### Leaf gas-exchange and fluorescence measurements

Thirty culms with the same flowering date were tagged at anthesis. Gas-exchange measurements were conducted at intervals of 7 days from anthesis to maturity. The leaf light-saturated net photosynthetic rate (*P_max_*), stomatal conductance (*g_s_*), and intercellular CO_2_ concentration (*C_i_*) of six tagged plants per plot were determined simultaneously. The average *P_max_* values of the six plants in each plot were taken as a replicate. *P_max_* was measured from 9:00 to 11:00 using a portable photosynthesis system (Li6400; LI-COR) at a light intensity of 1,200 μmol m^−2^ s^−1^. Leaf temperature during the measurements was maintained at 27.0 ± 0.1°C. The ambient CO_2_ concentration in the leaf chamber (*C_a-c_*) was adjusted as the atmospheric CO_2_ concentration (*C_a_*) (410 ± 1.5 μmol CO_2_ mol^−1^), and relative humidity was maintained at 60%. Data were recorded after equilibration to a steady state (~10 min). *PNUE* was calculated by dividing *P_max_* by *SLN*.

Steady-state fluorescence (*F_s_*), dark-adapted minimum fluorescence (*F_o_*), dark-adapted maximum fluorescence (*F_m_^′^*), and light-adapted maximum fluorescence (*F_m_*) were simultaneously measured using a portable fluorescent instrument (FMS-2, Hansatech, King’s Lynn, UK). Data were recorded after equilibration to a steady state. The maximum capture efficiency of excitation energy by open photosystem (PS)II reaction centres (*F_v_^′^/F_m_^′^*) and actual capture efficiency of excitation energy by open PSII reaction centres 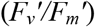 were estimated according to Genty *et al.* (1989).

### Measurement of mitochondrial respiration rate in the light (R_d_) and the CO_2_ compensation point related to C_i_ (Γ*)

*R_d_* and *Γ\** were measured by the following steps, which utilised the photorespiration rate being dependent on and *R_d_* being independent of photosynthetic photon flux density (*PPFD*, reviewed by Brooks and Farquhar, 1985; Bernacchi *et al.*, 2001). When the *P_max_*/*C_i_* response curves were prepared at a series of CO_2_ concentrations and at a battery of *PPFDs*, they intersected at one point where *P_max_* was the same at different *PPFDs.* Therefore, *P_max_* at that point represented –*R_d_*, and *C_i_* represented *Γ*.* In the present experiment, *R_d_* and *Γ\** were measured on different leaf layers from 0:00 h to 4:00 h (Brooks and Farquhar, 1985; Guo *et al.*, 2005, 2007). *PPFDs* were controlled as a series of 150, 300, and 600 μmol photons m^−2^ s^−1^. At each *PPFD, C_a-c_* was adjusted as a series of 25, 50, 80 and 100 μmol CO_2_ mol^−1^. The leaves were fixed in a leaf chamber with a *PPFD* of 600-μmol photons m^−2^ s^−1^ and a *C_a-c_* of 100-μmol CO_2_ mol^−1^ 30 min prior to initiating measurements.

### Stomatal density and stomatal aperture

Epidermal peels were stripped from leaves. Stomatal density was recorded under a microscope (Olympus Corp., Tokyo, Japan) in a 0.196-mm^2^ leaf area. A total of 1,000 stomatal apertures were measured under the microscope.

### Electron microscopy

Approximately 1–2-mm^2^ leaf sections were cut from the middle of each layer of leaves at anthesis using two razor blades, fixed in 2.5% glutaraldehyde (0.1 M phosphate buffer, pH 7.4), and post-fixed in 2% osmium tetroxide. Specimens were dehydrated in a graded acetone series and embedded in Epon 812. The leaf sections were cut on a Power Tome-XL ultramicrotome and stained with 2% uranyl acetate,. Then, cell wall thickness, chloroplast number, and chloroplast size were examined with an H-7650 transmission electron microscope.

### Calculation

#### Calculation of ETR

Total electron transport rate (*ETR*) was calculated from Eq. 1:

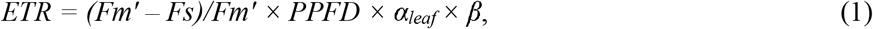

where *α_leaf_* is leaf absorbance, and *β* is the distribution of electrons between PSI and PSII. *α_leaf_* is dependent on chlorophyll content, and a curvilinear relationship between leaf absorption and chlorophyll content was observed by Evans (Evans *et al.*, 1996; Evans and Poorter, 2001). However, curvature was extremely low when chlorophyll content was >0.4 mmol m^−2^. According to Evans and Poorter (2001), the *α_leaf_* calculation demonstrates that *α_leaf_* is close to 0.85 (Asner *et al.*, 1998; Manter and Kerrigan, 2004). In this study, *α_leaf_* was also assumed to be 0.85, and *β* was assumed to be 0.5 (Ehleringer and Pearcy, 1983; Alvertssom, 2001).

#### Calculation of V_cmax_

The *V_cmax_* was calculated as described by Wilson *et al.* (2000).

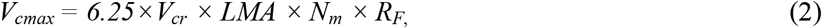

where 6.25 is the ratio of the weight of *Rubisco* to the weight of *N* in *Rubisco; V_cr_* is the specific activity of *Rubisco*, which is assumed to be only a function of temperature (20.7 μmol CO_2_ (g *Rubisco*)^−1^ s^−1^ at 25°C); *LMA* is leaf mass per unit area (g m^−2^); *N_m_* (g g^−1^) is the mass of *N* in the leaf per total mass of leaf; and *R_F_* is the apparent fraction of that *N* allocated to *Rubisco.*

#### Calculation of C_c_ and g_m_

Carbon dioxide concentration in chloroplasts (*C_c_*) and mesophyll conductance (*g_m_*) were calculated from Eqs. 6 and 7 (Harley *et al*., 1992; Epron *et al.*, 1995; Manter and Kerrigan, 2004):

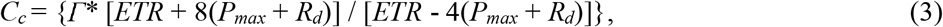

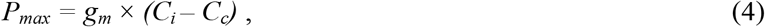

where *ETR* and *P_max_* were obtained from the gas-exchange and chlorophyll *a* fluorescence measurements conducted under saturating light; *R_d_* and *Γ\** were estimated as described above.

### Statistical analysis

Our results were analysed using DPS v 7.05 software (Hangzhou RuiFeng Information Technology Co., Ltd., Hangzhou, Zhejiang, China). Multiple comparisons were made after a preliminary F-test. Means were tested based on the least significant difference at *P* < 0.05.

## Results

### N allocation at the canopy level, plant N productivity, grain yield, and UTE

Over two wheat growing seasons, late-sown wheat plants accumulated less *N* per unit area than did that sown on the normal sowing date (Fig. 1). These reduced amounts of *N* were spread to a smaller plant population (Table 1) with higher *N* content in individual plants (Fig. 2). The above-ground biomass and *N* uptake (*AGN*) per unit area at anthesis were both reduced when the sowing date was delayed from 8 to 22 October (Fig. 1). As a result, similar plant *N* productivity was obtained from sowing to anthesis on both sowing dates. Late-sown wheat plants also accumulated less *AGN* from anthesis to harvest, but more biomass per unit area than did those with the normal sowing date. An average 30.8% increase in plant *N* productivity was obtained from anthesis to harvest under the later sowing date over the normal sowing date for the two wheat growing seasons (Fig. 1), indicating that improved plant *N* productivity with delayed sowing mainly resulted from more efficient *N* use during the post-anthesis period.

**Fig. 1.**
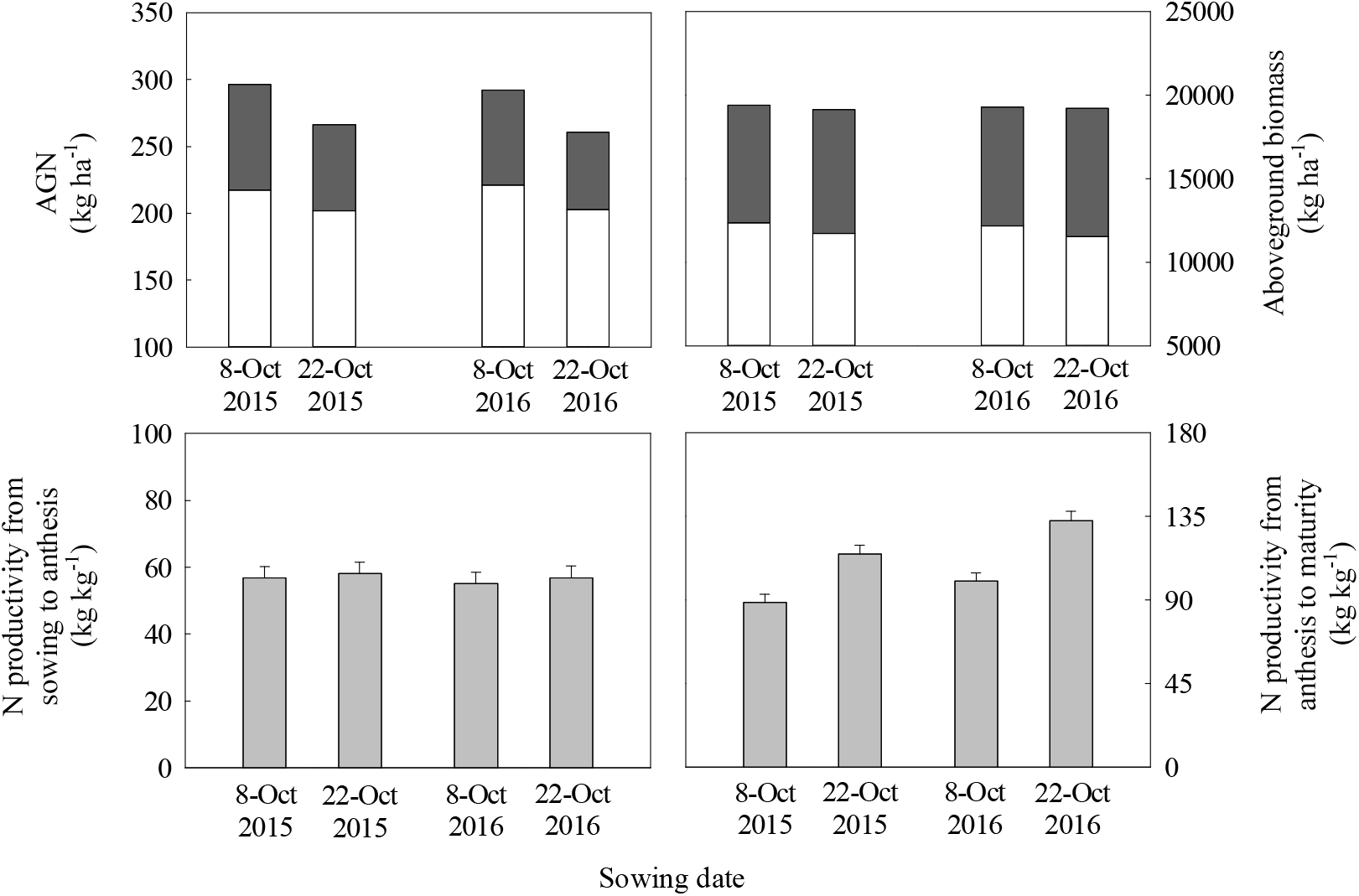
Aboveground *N* uptake (*AGN*) at anthesis (blank column) and maturity (blank column plus dark grey column), aboveground biomass at anthesis (blank column) and maturity (blank column plus dark grey column), *N* productivity from sowing to anthesis and from anthesis to maturity of winter wheat over two growing seasons. Vertical bars indicate standard error. Columns as follows: blank, aboveground AGN and biomass at anthesis; black, aboveground AGN and biomass from anthesis to maturity; light grey, *N* productivity.

**Fig. 2.**
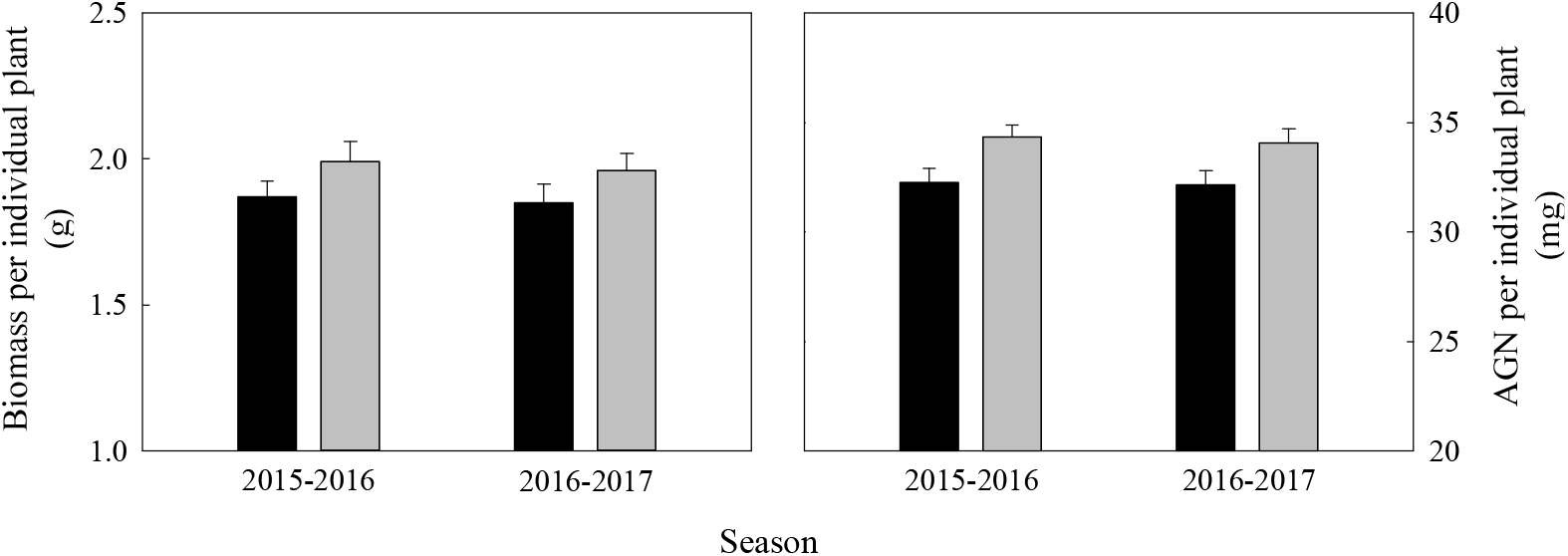
Biomass and aboveground nitrogen uptake (*AGN*) per individual plant at anthesis in winter wheat over two growing seasons. Vertical bars indicate standard errors. Columns as follows: black, 8 October; dark grey, 22 October.

**Table 1.** Grain yield, yield components, and nitrogen utilization efficiency (UTE) at harvest for two sowing dates over two wheat growing seasons. Values are means±standard errors of three replicates per treatment.

| Season | Sowing date | Grain yield (kg ha<sup>-1</sup>) | Spike number (10<sup>4</sup> ha<sup>-1</sup>) | Grain number per spike | Thousand grain weight (g) | UTE (kg kg<sup>-1</sup>) |
| --- | --- | --- | --- | --- | --- | --- |
| 2015-2016 | 8-Oct | 9316.7±232.9a | 662.4±16.6a | 37.2±0.93b | 39.5±0.94a | 31.4±0.85b |
|  | 22-Oct | 9243.7±225.1a | 590.1±14.7b | 41.8±1.1a | 39.4±1.0a | 34.7±0.92a |
| 2016-2017 | 8-Oct | 9432.8±243.8a | 670.5±16.8a | 37.6±1.0b | 39.2±1.0a | 32.3±0.81b |
|  | 22-Oct | 9378.6±238.6a | 605.7±15.3b | 41.6±1.2a | 39.6±1.1a | 36.0±0.86a |

Values followed by the same letter within a column and the same season are not significantly different at $P < 0.05$ as determined by the LSD test.

A high grain yield of >9,000 kg ha^−1^ was maintained, and *UTE* at harvest increased significantly when the sowing date was delayed (Table 1). In general, spike number per unit area decreased and spike grain weight increased as a result of increased spike grain number and unchanged grain weight (Table 1). These results suggest that trade-offs between spike number per unit area and grain number per spike resulted in similar grain yields between the two sowing dates.

### N allocation at the levels of the whole-plant and leaf, LMA, SLN, P_max_, and PNUE

With increased *AGN* and biomass of individual plants (Fig. 2), later-sown wheat plant allocated higher fractions of *AGN* and biomass at anthesis to the upper leaves, including the flag leaves and second leaves. The fraction of *AGN* and biomass allocated to lower-position leaves, including leaves 3 and 4, remained constant or decreased (Fig. 3). Different responses to delayed sowing in *LMA* and *SLN* were observed with leaf position in the canopy, as the area of all positioned leaves was not affected (Fig. 4). The *LMA* and *SLN* of the upper leaves increased, and those of lower positioned leaves remained constant or decreased (Fig. 5). The *N_m_* of all leaves remained unchanged. Therefore, changes in *SLN* were almost completely dependent on *LMA*.

**Fig. 3.**
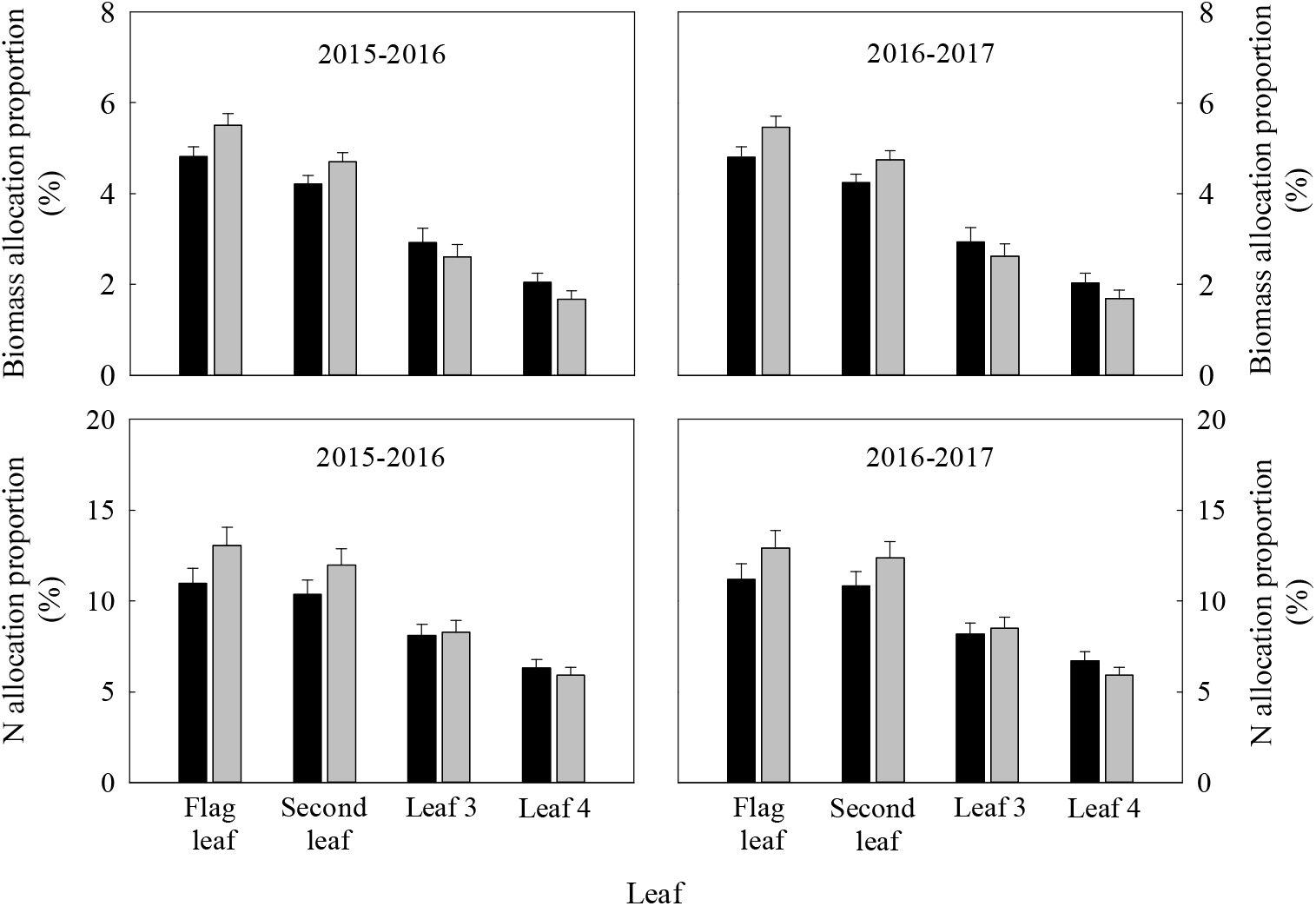
Biomass and *N* allocation proportion to flag leaf, second leaf, leaf 3, and leaf 4 in per individual winter wheat plant at anthesis over two growing seasons. Vertical bars indicate standard error. Columns as follows: black, 8 October; dark grey, 22 October.

**Fig. 4.**
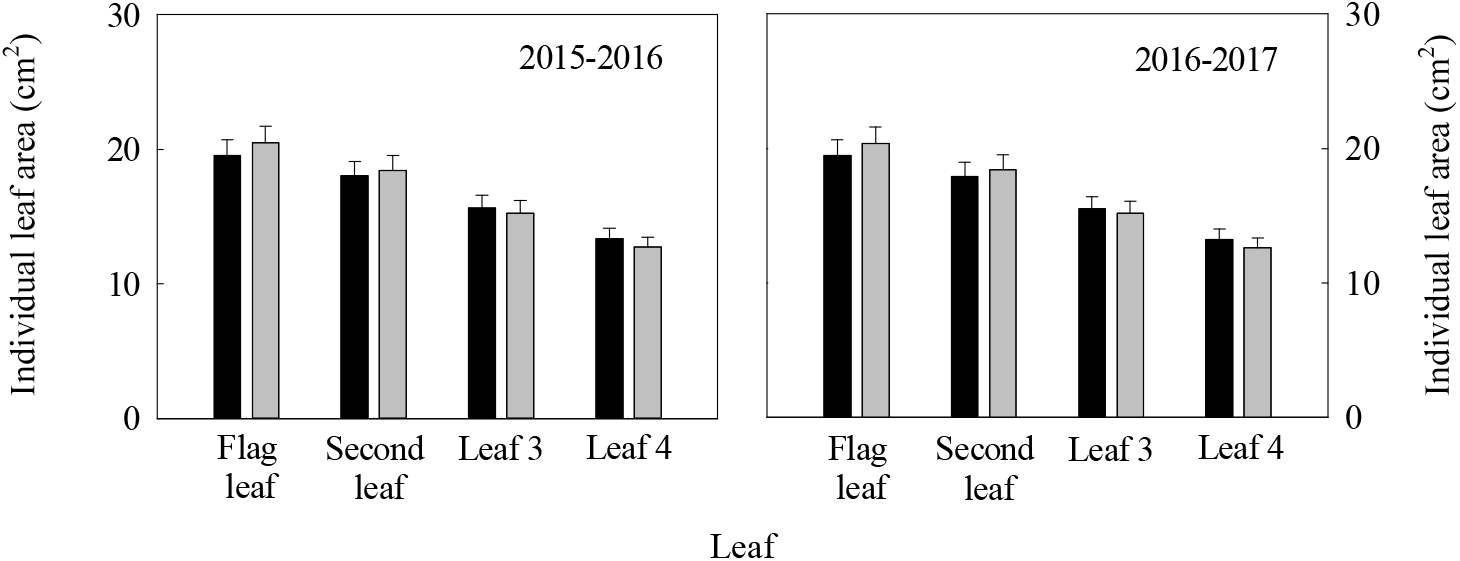
Area per individual leaf at different positions during anthesis in winter wheat over two growing seasons. Vertical bars indicate standard error. Columns as follows: black, 8 October; dark grey, 22 October.

**Fig. 5.**
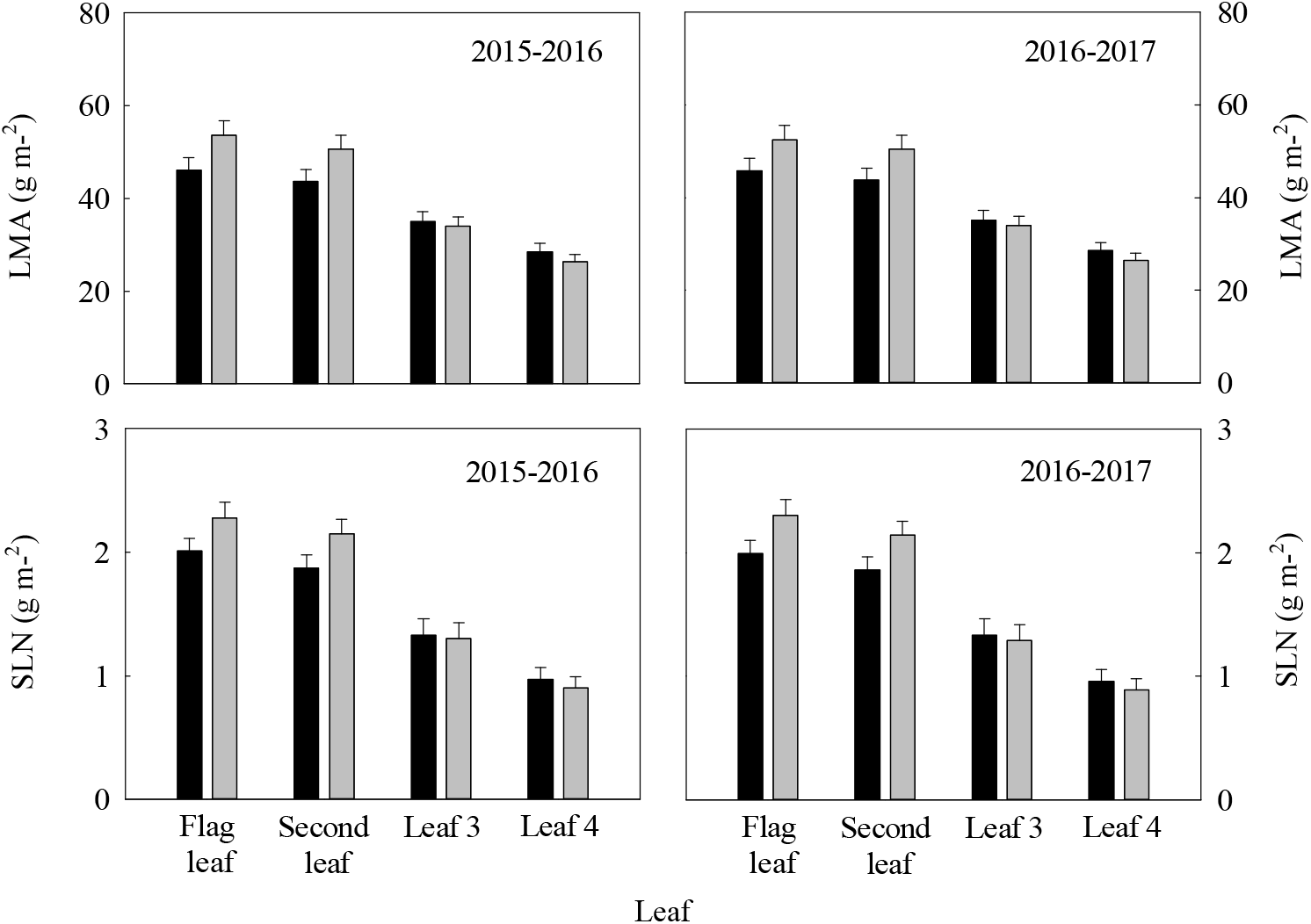
Leaf mass per area (*LMA*) and specific leaf nitrogen content (*SLN*) of different leaf layers at anthesis in winter wheat over two growing seasons. Vertical bars indicate standard error. Columns as follows: black, 8 October; dark grey, 22 October.

Improvement in *P_max_* and *PNUE* was attained in all leaves with delayed sowing. When the sowing date was delayed from 8 to 22 October, *P_max_* at anthesis increased over two growing seasons by, on average, 21.5%, 30.6%, 14.5%, and 25.4% in flag leaves, second leaves, and leaves 3 and 4, respectively (Fig. 6). Overall mean *PNUE* values at anthesis over the two growing seasons increased by 18.5%, 16.1%, 20.9%, and 31.2% in flag leaves, second leaves, leaf 3, and leaf 4, respectively (Fig. 6). The *PNUE* performance of the post-anthesis stage in different leaf layers was similar to that of anthesis (Supplemental Fig. 1).

**Fig. 6.**
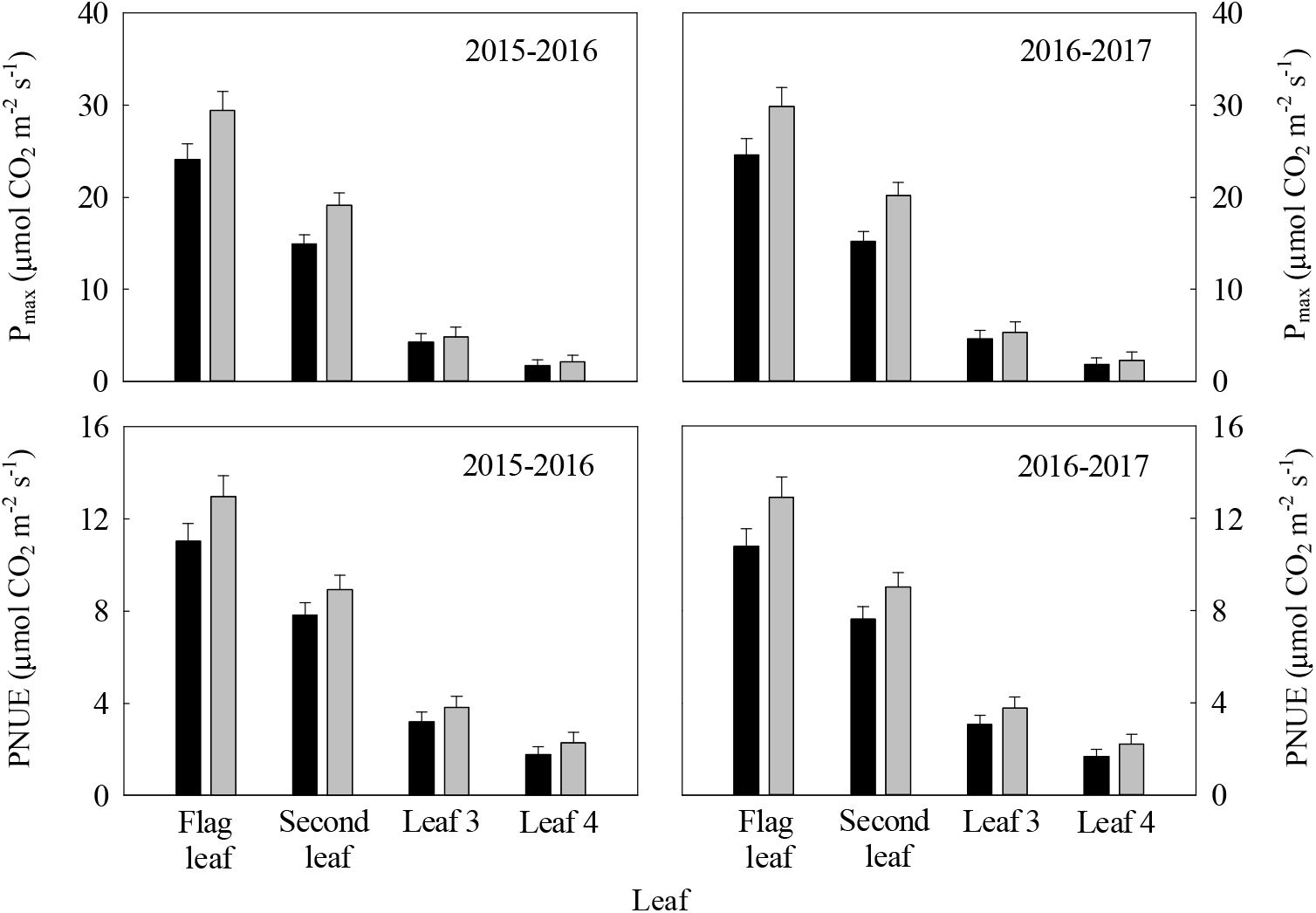
Light-saturated net photosynthetic rate (*P_max_*) and photosynthetic nitrogen use efficiency (*PNUE*) of different leaf layers at anthesis in winter wheat over two growing seasons. Vertical bars indicate standard error. Columns as follows: black, 8 October; dark grey, 22 October.

Taken together, these results suggest that strategies underlying improvements in *PNUE* in flag leaves and second leaves differed from that in leaves 3 and 4. Increased *P_max_* coupled with higher *LMA* and *SLN* in flag leaves and second leaves contributed to improve *PNUE*, whereas the combination of constant or reduced *SLN* and enhanced photosynthetic capability in leaves 3 and 4 resulted in improved *PNUE.*

### N allocation at the cellular level, Rubisco catalytic properties, and CO_2_ diffusion capacity

Optimising the functionality of *Rubisco* has large implications for improved plant productivity and resource use efficiency. Position-specific changes in transcript levels of the *mRNAs* coding *Rubisco* (Fig. 7) and the amount of *Rubisco* expressed as biomass and *N* content on a unit leaf area basis (Fig. 8) were observed with delayed sowing. As the allocation proportion of biomass and *N* to the cell wall decreased, the biomass and *N* content in the cell wall on a unit leaf area basis decreased for all positioned leaves (Fig. 8). The proportion of biomass and *N* allocated to total *Rubisco* and activated *Rubisco* in the flag and second leaves increased, while those in leaves 3 and 4 remained unchanged or decreased after the sowing date was delayed (Fig. 9). The *V_cmax_* of the upper leaves increased, and that in the lower leaves remained constant or decreased (Fig. 10).

**Fig. 7.**
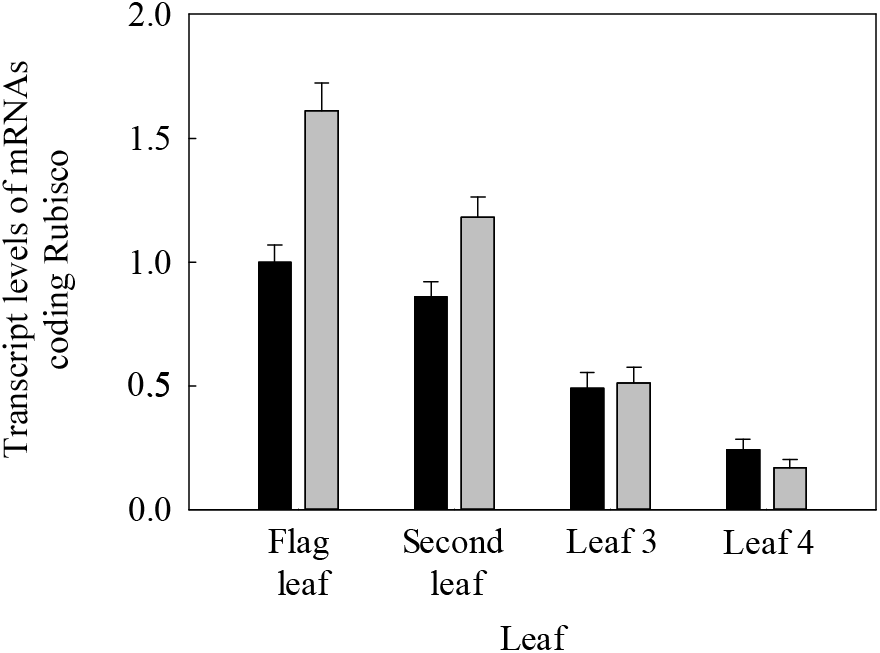
Transcript levels of *mRNAs* coding Rubisco per unit leaf area of different leaf positions at anthesis in winter wheat during the 2016-2017 season. Vertical bars indicate standard error. Columns as follows: black, 8 October; dark grey, 22 October.

**Fig. 8.**
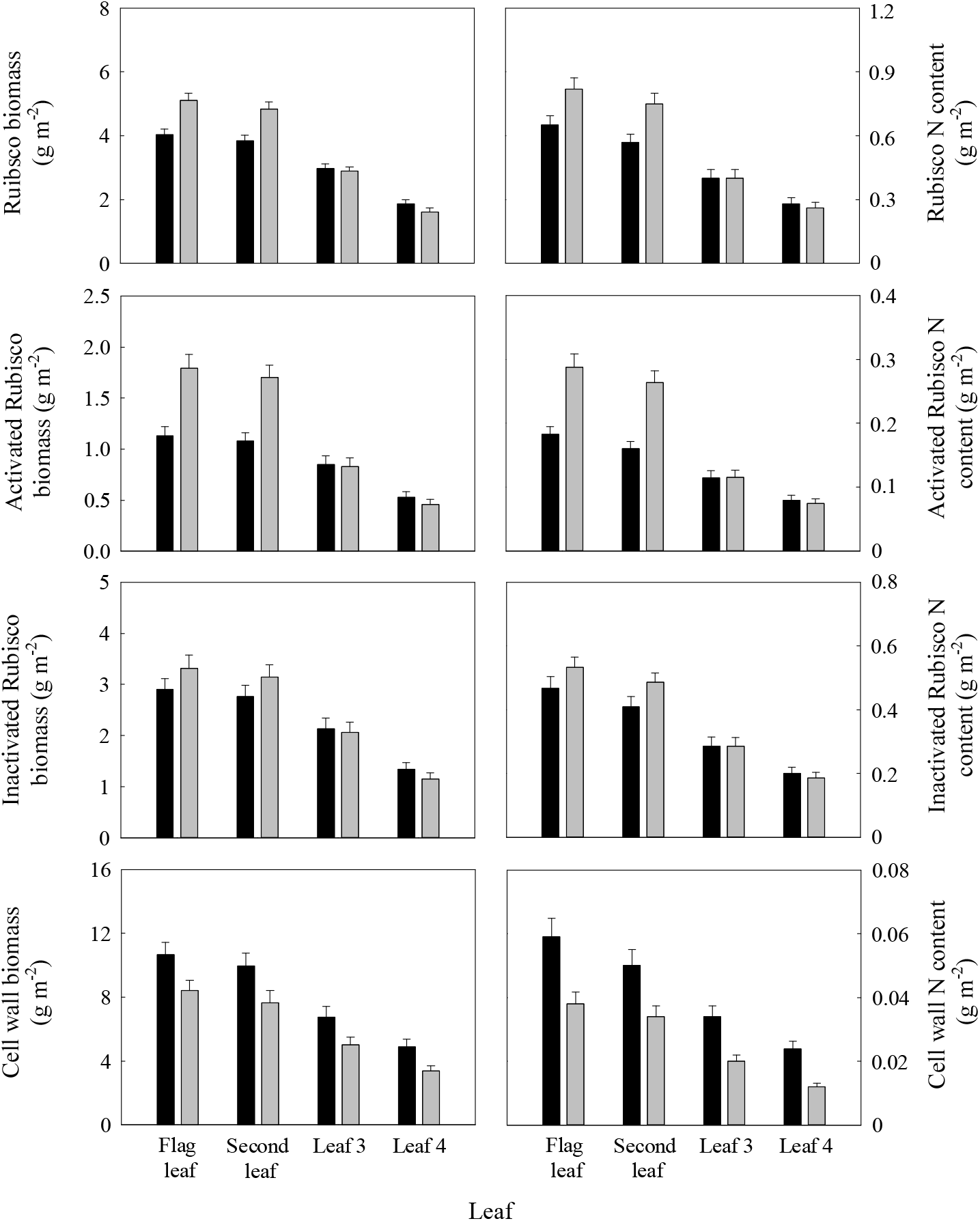
Rubisco biomass and *N* content, cell wall biomass and *N* content, activated Rubisco biomass and *N* content, and inactivated Rubisco biomass and *N* content per unit leaf area at different leaf positions during anthesis of winter wheat in the 2016-2017 season. Vertical bars indicate standard error. Columns as follows: black, 8 October; dark grey, 22 October.

**Fig. 9.**
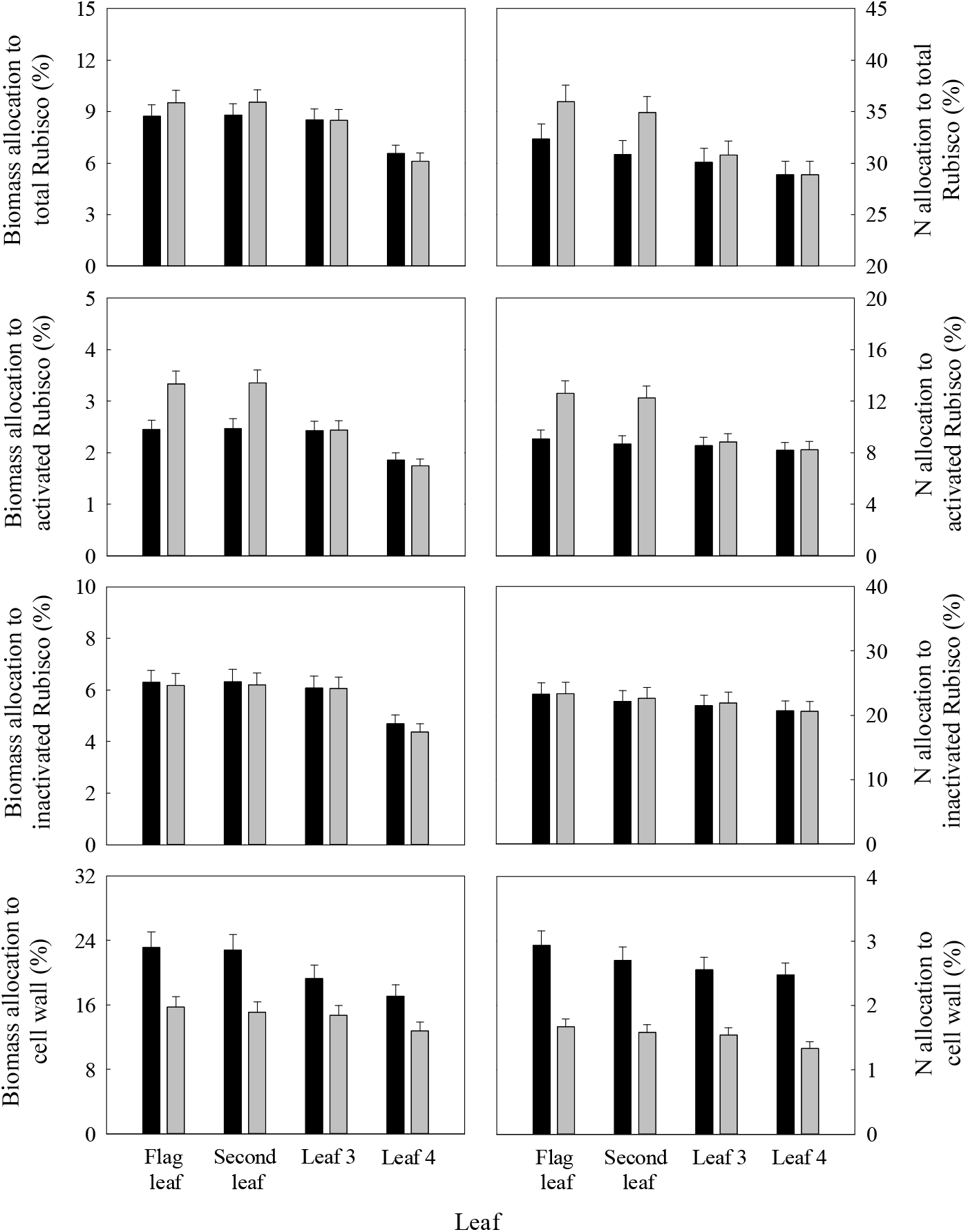
Biomass allocation to total Rubisco, *N* allocation to total Rubisco, biomass allocation to cell wall, *N* allocation to cell wall, biomass allocation to activated Rubisco, *N* allocation to activated Rubisco, biomass allocation to inactivated Rubisco, and *N* allocation to inactivated Rubisco per unit leaf area of different leaf positions at anthesis of winter wheat during the 2016-2017 season. Vertical bars indicate standard error. Columns as follows: black, 8 October; dark grey, 22 October.

**Fig. 10.**
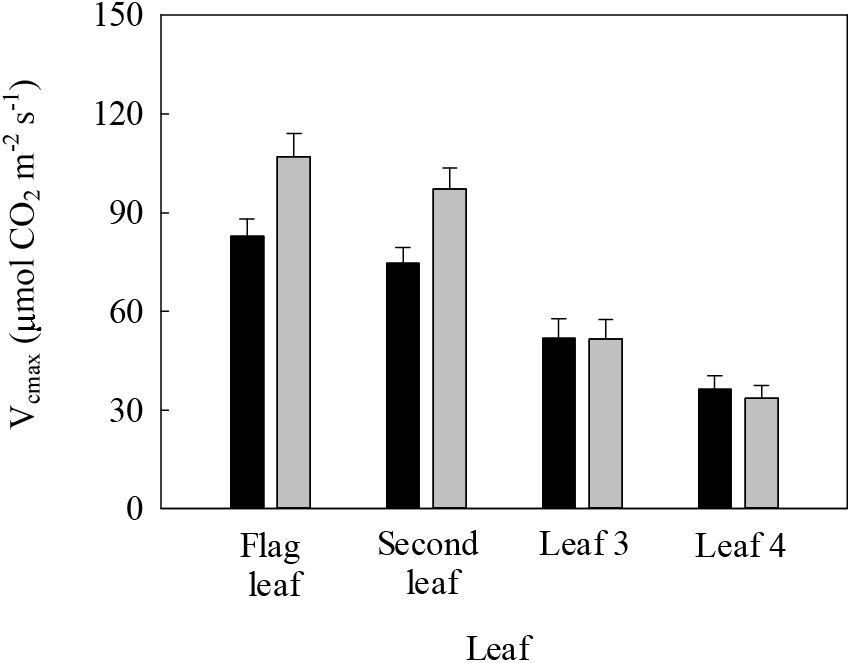
Maximum carboxylation rate limited by Rubisco (*V_cmax_*) per unit leaf area at different leaf positions during anthesis of winter wheat in the 2016-2017 season. Vertical bars indicate standard error. Columns as follows: black, 8 October; dark grey, 22 October.

Diffusional conductance of CO_2_ is the diffusive physiological determinant for the CO_2_ concentration at the *Rubisco* carboxylation site that directly affects net photosynthetic rate by limiting the amount of substrate (CO_2_) for fixation. The *g_s_, g_m_*, and associated traits, such as the number of stomata per unit area, stomatal aperture, cell wall thickness, chloroplast number, intercellular CO_2_ concentration, and chloroplast CO_2_ concentration, were measured or estimated. Higher *g_s_* values were obtained in leaves at all positions with later sowing compared with normal sowing, resulting in higher intercellular CO_2_ concentration (*C_i_*) (Fig. 11), which was associated with increased stomatal number per unit area (Fig. 12) and unchanged stomatal aperture. The *g_m_* values in all leaves at all positions were also enhanced by delayed sowing; consequently, higher chloroplast CO_2_ concentration (*C_c_*) was obtained (Fig. 11). The main reasons for this boosted *g_m_* include decreased cell wall thickness (Figs. 13, 14), increased chloroplast number per unit leaf area (Fig. 13), and unchanged chloroplast size in response to delayed sowing.

**Fig. 11.**
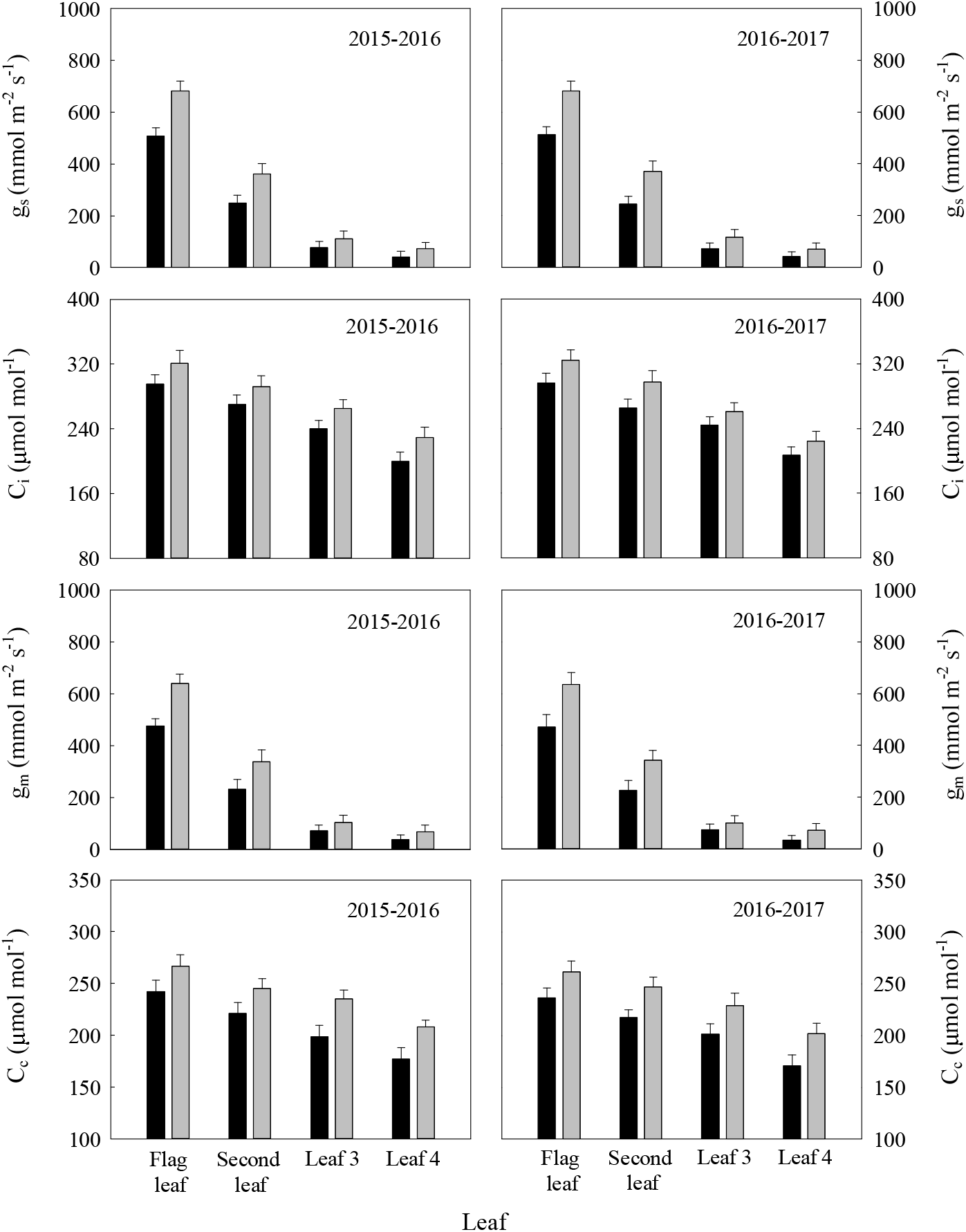
Stomatal conductance (*g_s_*), intercellular CO_2_ concentration (*C_i_*), mesophyll conductance (*g_m_*), and choroplastic CO_2_ concentration (*C_c_*) per unit leaf area at different leaf positions during anthesis in winter wheat over two growing seasons. Vertical bars indicate standard error. Columns as follows: black, 8 October; dark grey, 22 October.

**Fig. 12.**
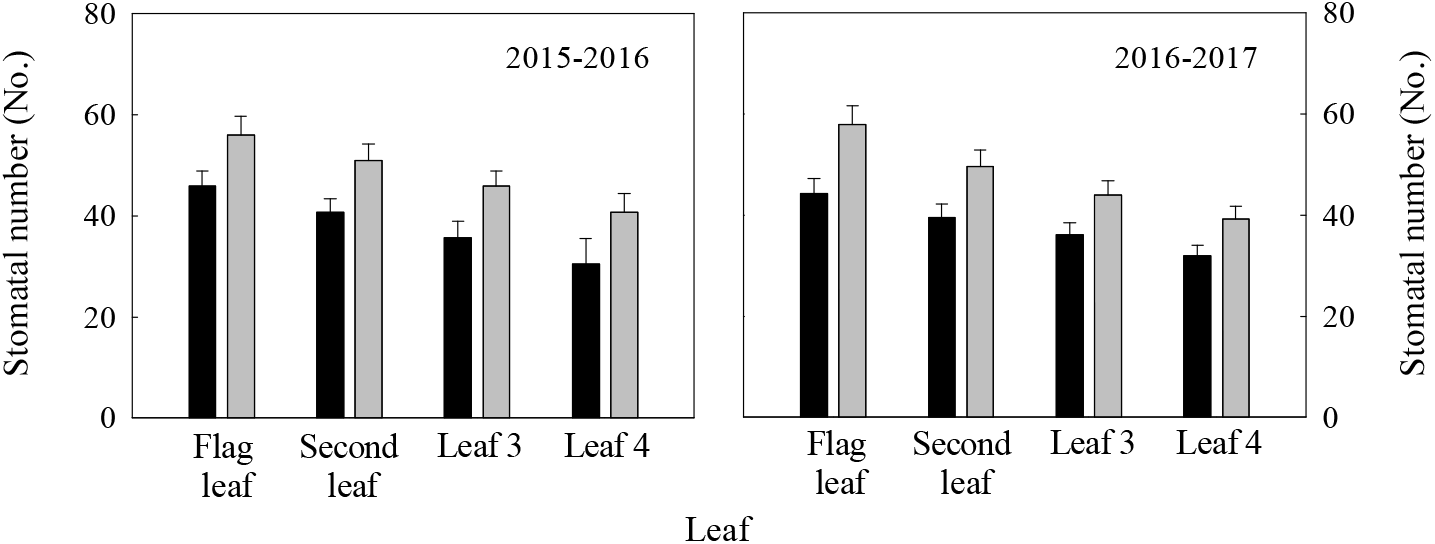
Stomatal number per unit leaf area at different leaf positions during anthesis in winter wheat over two growing seasons. Vertical bars indicate standard error. Columns as follows: black, 8 October; dark grey, 22 October.

**Fig. 13.**
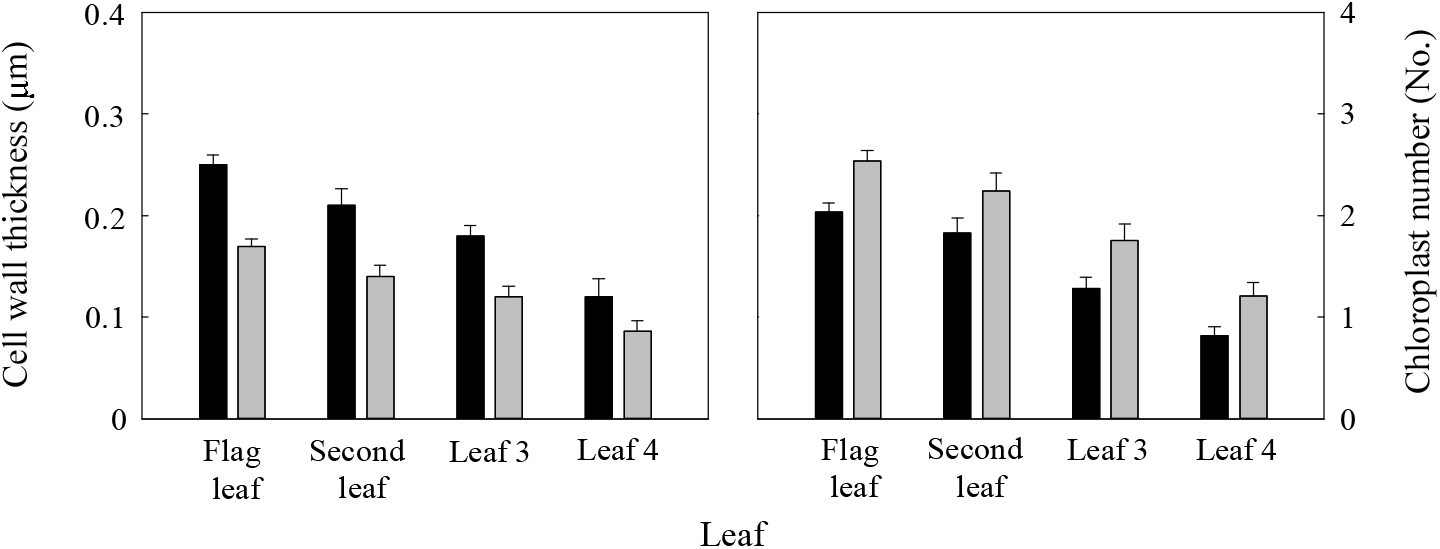
Cell wall thickness and chloroplast number per unit leaf area at different leaf positions during anthesis of winter wheat in the 2016-2017 season. Vertical bars indicate standard error. Columns as follows: black, 8 October; dark grey, 22 October.

**Fig. 14.**
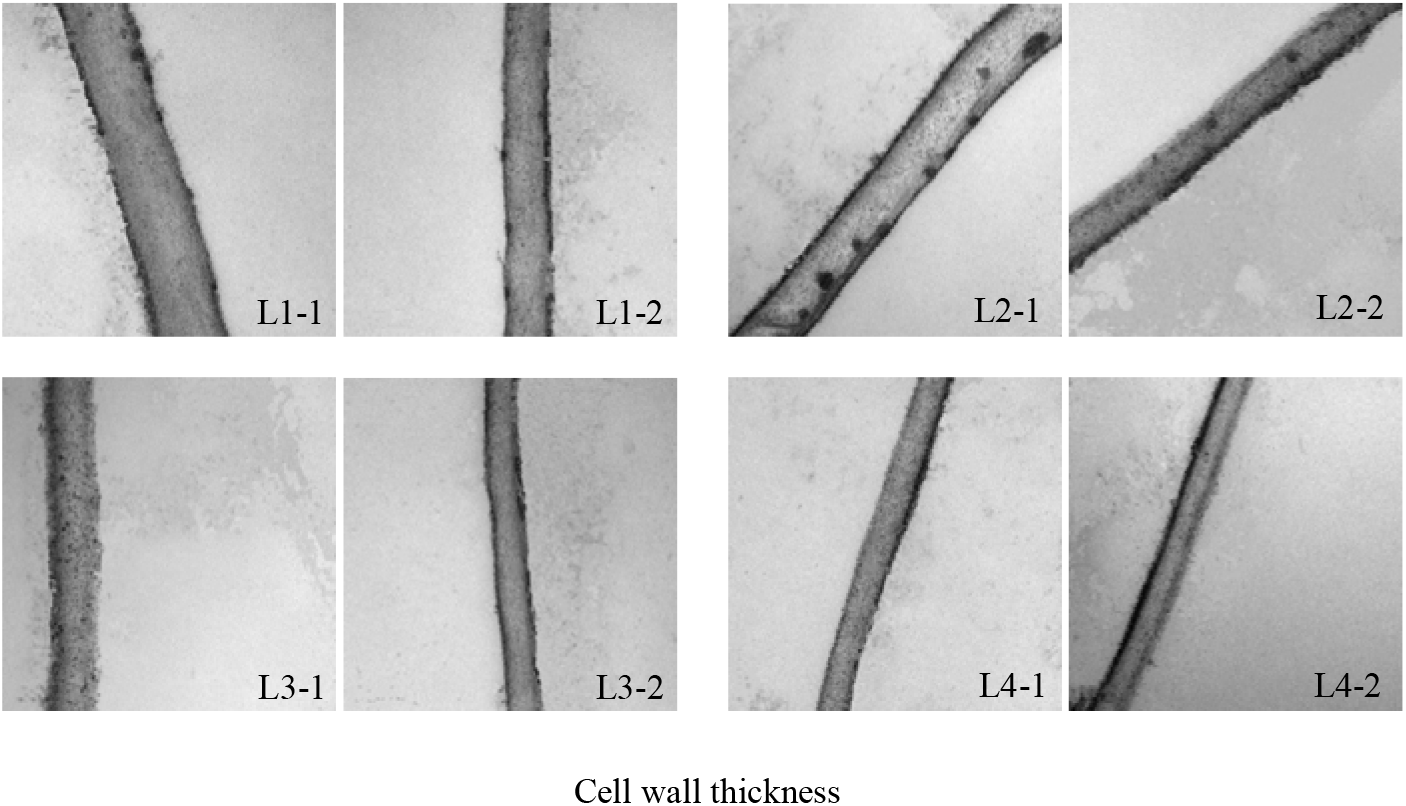
Estimates of cell wall thickness in winter wheat leaf (L1-1, flag leaf on 8 October; L1-2, flag leaf on 22 October; L2-1, second leaf on 8 October; L2-2, second leaf on 22 October; L3-1, leaf 3 on 8 October; L3-2, leaf 3 on 22 October; L4-1, leaf 4 on 8 October; L4-2, leaf 4 on 22 October) with normal and late sowing at anthesis by electron microscopy in the 2016-2017 season. All pictures are magnified 100,000×.

## Discussion

Manipulating *PNUE* at the leaf or whole-plant level will only be beneficial if it confers an improvement at the crop canopy level. As shown by Townsend *et al.* (2017), there is an opportunity to improve *PNUE* in the wheat canopy with no detriment to carbon gain or grain protein content by reducing the level of canopy *N.* In the present study, reduced canopy *AGN* at anthesis and at harvest were observed in response to delayed sowing. Later-sown wheat plants produced more biomass and grain yield on a unit area basis from anthesis to harvest with less *N* consumption than did those sown at a normal date, resulting in improved *PNUE* at the whole-plant level and *UTE* taking a reduced number of plants per unit area into consideration. As reduced total crop leaf area resulting from fewer plants per unit area was compensated for by enhanced photosynthetic capacity at the leaf and whole-plant levels, an improvement in *PNUE* at the crop canopy level was obtained while high grain yield was maintained.

It has long been recognised that the upper leaves serve as a major contributor to photoassimilates in the wheat grain (Waters *et al.*, 1980; Simpson *et al.*, 1983; Lopes *et al.*, 2006), while lower leaves contribute relatively little to grain yield during the grain-filling stage. Individual leaves require progressively less *N* from the top to the bottom of a canopy to maximise carbon assimilation (Gastal and Lemaire, 2002). Thus, an optimal correlation between the distribution of photosynthetic capacity, light, and *SLN* in flag leaves and second leaves is the main target for gains in yield potential, whereas leaves 3 and 4 are the main targets for gains in *PNUE* (Townsend *et al.*, 2018). In the present study, *AGN* at anthesis increased in individual plants due to a reduced number of plants per unit area. Leaf position-specific changes in *N* allocation were observed. *N* allocation to the upper leaves, such as the flag leaf and second leaf, increased, while *N* allocation to lower leaves, such as leaves 3 and 4, remained unchanged or decreased.

Canopy-level *PNUE* is a complex trait involving many plant characteristics and processes from leaf anatomy and composition to leaf physiology. Earlier studies concluded that increasing *PNUE* without considering grain yield required de-coupling of photosynthetic capacity and *SLN.* Strategies to improve *PNUE* while maintaining or increasing yield are lacking. This could potentially be achieved when *P_max_* is improved more than *SLN*.

Leaf conductance of CO_2_ and *Rubisco* kinetic parameters play key roles in carbon assimilation that are necessary for a proper understanding of photosynthetic performance under field conditions. High photosynthetic efficiency intrinsically demands tight coordination between traits related to CO_2_ diffusion capacity and leaf biochemistry. *V_cmax_* is the measure of the process by which *Rubisco* catalyses ribulose- 1,5-bisphosphate (*RuBP*) with CO_2_ to produce the carbon compounds that eventually become triose phosphates (e.g. glyceraldehyde-3P), the building block for sugars and starches. According to Wilson *et al.* (2000), variations in *V_cmax_* can be explained by changes in *LMA, N_m_*, and the *N* allocated proportion to *Rubisco* (*R_F_*). In the present study, *N_m_* remained unchanged in all leaves. The *LMA* and *R_F_* values in the flag and second leaves increased in response to delayed sowing, resulting in improved *V_cmax_*- However, *LMA* decreased in leaves 3 and 4, while *R_F_* remained unchanged in leaves 3 and 4, leading to unchanged *V_cmax_* in leaf 3 and a decrease in *V_cmax_* in leaf 4. The parallel increase between *LMA* and *R_F_* disagrees with a previous observation in which smaller *N* partitioning into *Rubisco* was observed against larger *N* partitioning into cell walls with increasing *LMA* (Poorter and Evans, 1998; Onoda *et al.*, 2004; Takashima *et al.*, 2004; Wright *et al.*, 2005; Harrison *et al.*, 2009; Hidaka *et al.*, 2009). The main reason for this difference may be related to whether interspecific (previous study) or intraspecific comparisons were made (present study).

As *V_cmax_* represents the maximum carboxylation rate under both light-saturated and CO_2_-saturated conditions, *P_max_* was measured under light saturation but at a normal ambient CO_2_ concentration; therefore, the difference between the two parameters reflected a limitation on photosynthetic capacity exerted by the CO_2_ supply.

The *g_m_* value, a limiting factor for CO_2_ diffusion to carboxylation sites in the stroma, is usually tightly coregulated with *g_s_* (Flexas *et al.*, 2013). The *g_m_* value depends on the surface area of mesophyll cells exposed to the intercellular air space and the thickness of the mesophyll cell walls (Evans *et al.*, 1994, 2009; Tholen and Zhu, 2011; Tosens *et al.*, 2012). The finding that *g_m_* is constrained by large *LMA* has been reviewed previously (Flexas *et al*., 2008), and the underlying reason is mostly related to the thicker cell walls observed in species with high *LMA*, which significantly limits CO_2_ diffusion inside leaves (Parkhurst, 1994; Hanba *et al.*, 1999; Wright *et al.*, 2005; Hidaka *et al.*, 2009; Peguero-Pina *et al.*, 2012; Tosens *et al.*, 2012; Tomás *et al.*, 2013). In contrast to previous reports, thinner cell walls in leaves at all positions were observed with larger *LMA* due to reduced biomass allocation to the cell wall under the delayed sowing condition. Moreover, an increase in the number of chloroplasts per unit leaf area allowed for a larger surface area of mesophyll cells exposed to intercellular air space. Higher *g_s_* values were obtained in leaves at all positions on plants sowed later due to an increased number of stomata per unit area and an unchanged stomatal aperture, which is also helpful for increasing chloroplastic CO_2_ concentration (*C_c_*). Combining these observations, we propose that the dominant mechanism for improved *P_max_* in lower leaves in the canopy is enhanced CO_2_ diffusion capacity, and that in the upper leaves is dependent on the combination of *Rubisco* catalytic properties and CO_2_ diffusion capacity.

## Conclusion

Optimal *N* allocation was achieved at several integration levels in response to delayed sowing. A limited amount of *N* was spread over a reduced plant population at the crop-canopy level, which in turn resulted in increased *N* content in individual plants. An increased fraction of *N* was allocated to upper leaves, including the flag and second leaves, which are main contributors of photoassimilates to grain filling. A decreased fraction of *N* was allocated to lower leaves, including leaves 3 and 4, which contribute relatively little to grain yield during grain filling. At the leaf level, *N_m_* was constant between the two sowing dates. The *LMA* of upper leaves increased as a result of investing more biomass in a given area. As *N_m_* remained constant, the *SLN* of these leaves increased, whereas *LMA* and *SLN* of the lower leaves remained unchanged or decreased. At the cellular level, larger proportions of *N* were allocated to *Rubisco* (both total and activated), which alone or along with increased *LMA* increased *V_cmax_* in the upper leaves, while it remained unchanged or decreased in the lower leaves. Higher *g_s_* values were obtained in leaves at all positions with later sowing due to the increased number of stomata per unit area and unchanged stomatal aperture, which is helpful for increasing *C_i_*. Thinner cell walls and an increased number of chloroplasts per unit leaf area allowed for increases in *g_m_* and *C_c_*. Tight coordination between *Rubisco* catalytic properties and CO_2_ diffusion capacity led to improved *P_max_* in the upper leaves, whereas improvement in *P_max_* in lower leaves is dependent on enhanced CO_2_ diffusion capacity. Outperformance by *P_max_* over *SLN* led to improved *PNUE* in upper leaves. Enhanced *P_max_* coupled with unchanged or decreased *SLN* resulted in improved *PNUE* in lower leaves. In summary, optimal *N* allocation accounted for the improvement in *PNUE* at the crop-canopy level while maintaining a high grain yield.

## Acknowledge

This work was supported by the Chinese National Basic Research Program (2015CB150404) and Funds of Shandong “Double Top” Program (SYL2017YSTD05).

## Abbreviations

*C_c_*: chloroplastic CO_2_ concentration
*C_i_*: intercellular CO_2_ concentration
*ETR*: total electron transport rate
*g_m_*: mesophyll conductance
*g_s_*: stomatal conductance
*J_max_*: light-saturated potential rate of electron transport
*LMA*: leaf mass per area
*N_m_*: the mass of nitrogen in the leaf per total mass of leaf
*P_max_*: light-saturated net photosynthetic rate
*PPFD*: photosynthetic photon flux density
*PNUE*: photosynthetic nitrogen-use efficiency
*Rubisco*: ribulose-1, 5-bisphosphate carboxylase/oxygenase
*R_f_*: nitrogen allocated to *Rubisco*
*SLN*: specific green leaf area nitrogen
*V_cmax_*: maximum carboxylation rate

